# Evolution of Resistance Against CRISPR/Cas9 Gene Drive

**DOI:** 10.1101/058438

**Authors:** Robert L. Unckless, Andrew G. Clark, Philipp W. Messer

## Abstract

The idea of driving genetically modified alleles to fixation in a population has fascinated scientists for over 40 years^1,2^. Potential applications are broad and ambitious, including the eradication of disease vectors, the control of pest species, and the preservation of endangered species from extinction^3^. Until recently, these possibilities have remained largely abstract due to the lack of an effective drive mechanism. CRISPR/Cas9 gene drive (CGD) now promise a highly adaptable approach for driving even deleterious alleles to high population frequency, and this approach was recently shown to be effective in small laboratory populations of insects^4–7^. However, it remains unclear whether CGD will also work in large natural populations in the face of potential resistance mechanisms. Here we show that resistance against CGD will inevitably evolve unless populations are small and repair of CGD-induced cleavage via nonhomologous end joining (NHEJ) can be effectively suppressed, or resistance costs are on par with those of the driver. We specifically calculate the probability that resistance evolves from variants at the target site that are not recognized by the driver's guide RNA, either because they are already present when the driver allele is introduced, arise by *de novo* mutation, or are created by the driver itself when NHEJ introduces mutations at the target site. Our results shed light on strategies that could facilitate the engineering of a successful drive by lowering resistance potential, as well as strategies that could promote resistance as a possible mechanism for controlling a drive. This study highlights the need for careful modeling of CGD prior to the actual release of a driver construct into the wild.

---

CGD involves the design of an autonomous genetic construct that contains Cas9, a guide RNA (gRNA) targeting a specific site in the genome, and flanking homology arms to facilitate incorporation into that genome^3,4^. In heterozygotes, the driver can cleave the sister chromosome at the target site and insert itself via homology-directed repair, converting heterozygotes for the driver construct into homozygotes (Fig. 1a). In addition, arbitrary genetic segments can be included in the construct that will be transmitted alongside the driver.

**Figure 1.**
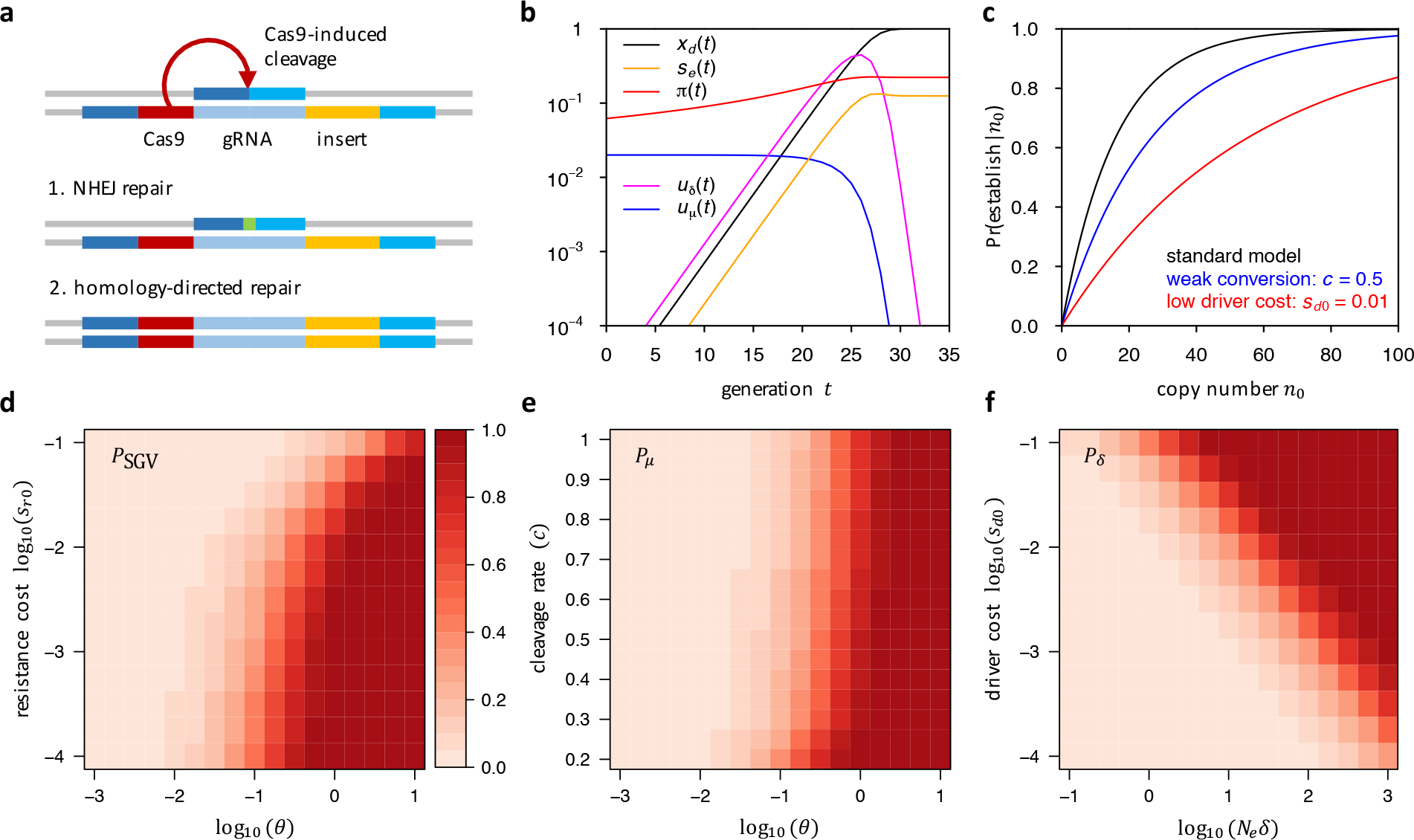
CGD dynamics and resistance probabilities. **a**, In heterozygotes, the driver cleaves the wildtype chromosome at the target site. Cleavage-repair by NHEJ can create a resistance allele, e.g., by introducing an indel (green) at the target site, whereas homology-directed repair will create a driver homozygote, **b**, The driver initially grows exponentially and fixes after ~30 generations in our standard model; *u_μ_(t)* is proportional to the frequency of wildtypes and thus decreases as the driver becomes more frequent; *u_㊴_(T)* is proportional to the frequency of driver/wildtype heterozygotes and is thus maximal for intermediate driver frequencies; *π(t)* increases with driver frequency, but is already quite high in generation zero, because resistance alleles from the SGV only have to survive a few generations of drift before their relative selective advantage becomes noticeable, **c**, Probability that resistance establishes from the SGV as a function of the initial copy number (*n_0_*) in which resistance alleles are present when the driver is introduced. Shown are the standard model, a scenario in which the driver cleaves at only 50% efficiency, and a scenario where driver cost is ten times smaller. **d**, *P*_SGV_ as a function of *θ* under codominant resistance cost. Higher resistance costs reduce *P*_SGV_. **e**, *P_μ_* as a function of *θ* and cleavage rate *c*. Higher *c* slightly increases *P_μ_*. **f**, *P_δ_* as a function of *θ* and driver cost. Higher driver cost decreases *P_δ_*. In **d**,**e**, we varied *N_e_* while keeping *μ* = 10^−8^ constant. In **f**, we varied *N_e_* while keeping *δ* = 10^−6^ constant.

As a proof of concept, Gantz and Bier^4^ showed that CGD was highly effective at spreading a mutant allele in a laboratory cross of *D. melanogaster*. A similar construct conferring resistance to *Plasmodium falciparum* was successfully introduced into a laboratory population of *Anopheles* mosquitoes^5,7^. Theoretical analyses in a Wright-Fisher framework with conversion, selection, and drift also showed that CGD can in principle lead to rapid fixation of a driver allele even when it carries a substantial fitness cost^8,9^.

The key to determining whether CGD can actually transform natural populations is in understanding the likelihood that resistance will evolve against the driver construct. Such resistance may arise from alleles at the target locus that do not drive themselves but also cannot be converted by the driver, because they are not recognized by the gRNA. Cas9 tolerates only a few mismatches between the 20 bp-long target sequence and gRNA, and none in the upstream PAM sequence^10^. It stands to reason that most indels inside the target sequence will prevent binding as well. Despite experimental evidence suggesting that such resistance can evolve quickly^11,12^, this problem has not been addressed in theoretical models of CGD.

To study the probability that resistance evolves against CGD we devised a population genetic model of a single locus with three classes of alleles: wildtype (*0*), driver (*d*), and resistance (*r*). The driver cleaves the wildtype allele in a driver/wildtype heterozygote at rate *c*. In a fraction *δ* of cases, repair of such breaks creates a resistance allele when NHEJ introduces an indel at the target site. Resistance alleles can also arise by *de novo* mutation in wildtype alleles at rate *μ* specifying an effective rate including all possible mutations that create a resistance allele. For simplicity, we consider resistance a binary trait — alleles are completely resistant or completely susceptible. We set the fitness of wildtype homozygotes to *ω_00_* = 1, whereas all other genotypes can carry arbitrary fitness cost, *ω_ij_* = 1−*s_ij_* ≤ 1.

In this model, we can express the expected values of allele frequencies *x_d_*(*t*), *x_r_*(*t*), and *x_o_*(*t*) as functions of their frequencies in the previous generation, the fitness costs of the different genotypes, and the rates *c*, *μ* and *δ* (SI section 1.2). These equations then fully describe the dynamics of allele frequencies from a given set of starting frequencies in a deterministic scenario, which will be appropriate once allele frequencies are large enough such that genetic drift can be ignored. Alternatively, these equations can be incorporated into a Wright-Fisher framework that explicitly takes drift into account.

We distinguish three mechanisms by which resistance alleles can originate: (i) standing genetic variation (SGV); (ii) *de novo* mutation after the driver is introduced (specified by i); and (iii) double strand break-repair by NHEJ (specified by *δ*). Let *P*_SGV_, *P_μ_* and *P_δ_* denote the probabilities that resistance evolves by each particular mechanism, assuming that it does not evolve by any of the other mechanisms. Each probability depends on the supply of resistance alleles by its mechanism and the probability *π*(*t*) that a resistance allele successfully establishes in the population (i.e., is not lost by drift) when it is initially present in a single copy in generation *t*.

Calculation of *π*(*t*) is complicated by the fact that the effective fitness advantage of a resistance allele over the population mean, s_e_(t), is a function of the driver frequency and thus is time-dependent. When the driver is at low frequency, *s_e_*(*t*) will be small or could even be negative if resistance alleles carry costs themselves. Only as the mean population fitness decreases when a deleterious driver spreads will *s_e_(t)* increase. We calculated *π*(*t*) in our model by adapting results from Uecker and Hermisson^13^ (SI section 2.1).

To quantify the expected supply of resistance alleles from SGV, we assumed mutation-selection-drift balance prior to the introduction of the driver, which is determined by the fitness costs of resistance alleles in the absence of a driver and the population-level mutation rate toward resistance alleles, *θ* = 4*N_e_μ* where *N_e_* is the variance effective population size^14^. *De novo* mutation creates resistance alleles at rate *u_μ_*(*t*) ≈ 2*Nμ*[(*1−x_d_*)^2^+*x_d_(1−x_d_*)(*1−c*)], where *N* is the census population size. This rate is proportional to the number of wildtype alleles and thus decreases as the driver spreads. NHEJ creates resistance alleles at rate *u_δ_*(*t*) ≈ 2*Nδcx_d_*(*l−x_d_*), which is proportional to the number of wildtype/driver heterozygotes and thus is highest when the driver is at intermediate frequency. Combining these rates with our analytic result for *π(t)* enabled us to calculate the individual resistance probabilities *P*_SGV_, *P_μ_* and *P_δ_*, provided in Equations (16), Equation (19), and Equation (22) in SI. Note that once a driver has fixed in the population, resistance can no longer evolve by any of the above mechanisms.

To dissect how resistance probabilities depend on the different parameters, we first defined a “standard” model and then varied its parameters independently. This model assumes efficient conversion, *c* = 0.9, and cost-free resistance, *s_r0_* = *s_rr_* = 0. Driver alleles, by contrast, carry a substantial cost that is codominant, *s_d0_* = *s_dd_/2* = 0.1. We set *δ* = 10^−6^, *μ* = 10^−8^, and assume *N* = *N_e_* = 10^6^. Driver alleles are introduced at frequency *x_d_*(0) = 10^−5^. Figure 1b shows how *x_d_*(*t*), *s_e_*(*t*), *π*(*t*), *u_μ_*(*t*), and *u_δ_*(*t*) change over time in this model when no resistance alleles are yet present.

Whether resistance evolves from SGV is primarily a function of the number of resistance alleles present when the driver is introduced (Fig. 1c), which is determined by *θ* in our standard model with cost-free resistance. If resistance alleles carry fitness costs themselves, this can lower *P*_SGV_ by reducing their expected frequency in the SGV (Fig. 1d). However, this effect becomes significant only when resistance costs are codominant, not when they are recessive (SI Fig. 2c,Fig 2d). Lower cleavage rates only slightly decrease establishment probabilities (Fig. 1c). This is because for a slower driver it will take longer until resistance alleles from the SGV experience their fitness advantage, increasing their chances of being lost to drift and/or selection in the meantime. Similarly, lower driver costs decrease *π*(*t*) and thus *P*_SGV_, as resistance alleles will have a smaller fitness advantage (SI Fig. 2b). Generally, as long as resistance alleles do not carry large costs themselves and provide any fitness advantage over the driver, evolution of resistance from the SGV is practically assured whenever *θ* > 0.1, whereas it remains unlikely for *θ* ⪡ 0.1 (Fig. 1d, SI Fig. 2), consistent with general results for the probability of adaptation from SGV after an environmental shift^14–17^.

**Figure 2.**
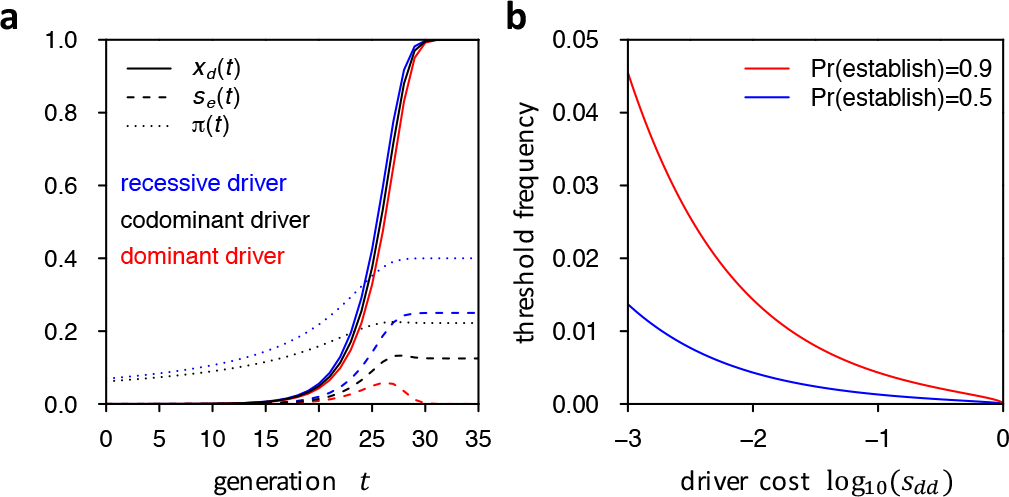
Recessive and dominant driver costs. **a**, Driver allele frequency trajectories, effective fitness advantage of resistance allele, and establishment probabilities under recessive, codominant, and completely dominant driver fitness costs in our standard model. *π(t)* is not visible in the dominant scenario as it is very close to zero. **b**, Threshold frequencies *x_0_* at which a resistance allele needs to be present to have the given establishment probability in the case of a completely dominant driver with cost *s_dd_.*

The likelihood that resistance evolves from *de novo* mutation also depends strongly on *θ*, with resistance becoming likely once *θ* > 1. Lower cleavage rates increase *P_μ_* (the opposite effect they have on *P*_SGV_) because a slower driver provides more time for resistance alleles to emerge before the driver can fix (Fig. 1e). Lower driver costs again reduce *P_μ_* by lowering *π*(*t*) (SI Fig. 3b). We find that in most scenarios *P*_SGV_ is higher than *P_μ_* Thus, resistance is more likely to evolve from alleles in the SGV than from *de novo* mutation after introduction of the driver, unless the drive is very slow.

**Figure 3.**
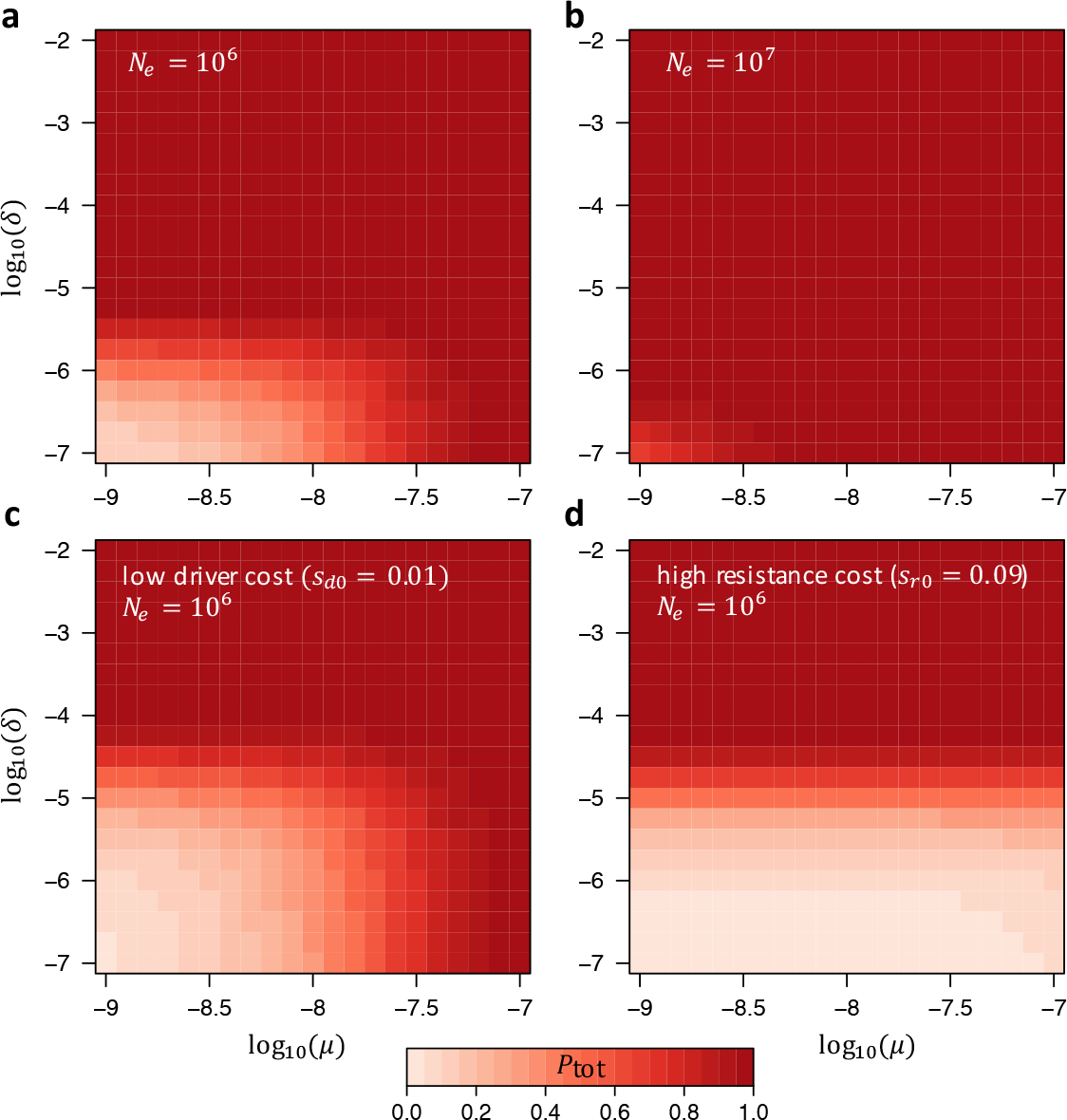
Overall probability that resistance evolves. **a**,**b**, *P*_tot_ as a function of *μ* and *δ* in our standard model for two different effective population sizes *N_e_* = 10^6^,10^7^. **c**, Scenario with low driver cost: *s_d0_* = *s_dd_*/2 = 0.01. This allows for somewhat larger values of *δ* before resistance evolves. The dependence on *μ*, however, does not change much, since *P*_SGV_ does not depend strongly on driver cost. **d**, Scenario in which resistance alleles carry a (codominant) fitness cost that is 90% that of the driver: *s_r0_* = 0.9*s_d0_* = *s_rr_*/2 = 0.09. In this case, resistance is unlikely to evolve from SGV or *de novo* mutation. However, it will still evolve by NHEJ whenever *δ* > 10^−5^.

The key parameter determining the likelihood of resistance from NHEJ is the product *N_e_δ*, similar to the role played by *θ* = 4*N_e_μ* for *de novo* mutation. Whenever *N_e_δ* > 1, resistance becomes likely (Fig. 1f). In contrast to *de novo* mutation, the cleavage rate here has very little impact on *P_δ_* (SI Fig. 4b). This is because the overall number of resistance alleles that arise by NHEJ depends only weakly on *c* (SI section 2.4). Lower fitness costs of the driver again decrease *P_δ_* by reducing *π*(*t*) (Fig. 1f).

**Figure 4.**
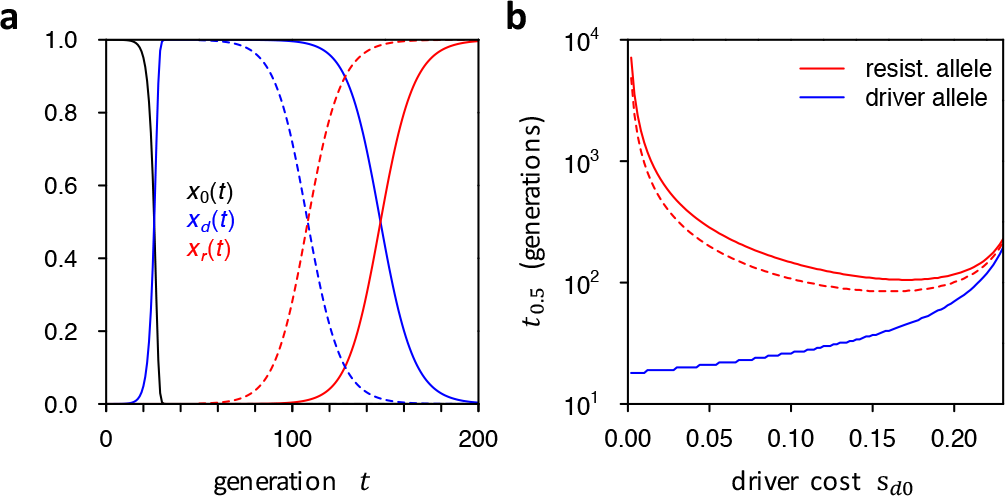
Replacement of driver by resistance allele. **a**, Deterministic frequency trajectory of driver allele in our standard model and its replacement by a resistance allele from the SGV that was initially present as a single copy (solid line) or one hundred copies (dashed line) when the driver was introduced. **b**, Time until resistance allele and driver allele reach 50% population-frequency in a when varying driver cost *s_d0_*. Once driver cost reaches *s_d0_* ≈ 0.25 in our model, the driver will be outpaced by the resistance allele before it can reach 50%.

So far, we have assumed that driver costs are codominant. Recessive driver costs will facilitate faster initial spread of the driver, whereas dominant costs will slow it down. However, these differences are only marginal for the selection coefficients and cleavage rate in our standard model (Fig. 2a), as frequency changes of the driver allele between generations are still dominated by conversion, rather than selection.

Dominance of the driver can nevertheless have strong impact on the probability that resistance arises, due to its effects on the fitness of driver/resistance heterozygotes. If driver costs are recessive, *s_e_(t)* will be larger than in the codominant case once the driver becomes more frequent, increasing *π*(*t*) (Fig. 2a). By contrast, if driver costs are completely dominant, driver/resistance heterozygotes will have no fitness advantage over driver homozygotes. Whether resistance can still evolve in this case is analogous to the question of whether a recessive beneficial mutation can establish, which requires a high initial frequency of that mutation so that homozygotes will be present in the population (Fig. 2b, SI section 3.2). For example, when the driver has a dominant fitness cost of *s_dd_* = 0.1, resistance alleles would have to be present at ~0.1% frequency in order to have a 50% chance of successfully establishing. Note that we would expect this to be the case in scenarios with *δ* > 10^−3^ (SI section 3.1).

Given *P*_SGV_, *P_μ_* and *P_δ_*, we can calculate the total probability that resistance evolves by any of the mechanisms: *P*_tot_ = 1−(1−*P*_SGV_)(1−*P_μ_*)(1−*P_δ_*). Figure 4 shows *P*_tot_ over a wide parameter space. Consistent with the above results, *P*_tot_ depends primarily on three factors: *N_e_*, *μ*, and *δ*. Resistance is generally likely to evolve whenever either *θ* > 0.1, or *N_e_δ* > 1, as long as resistance alleles provide any fitness advantage over the driver and driver costs are not completely dominant.

We expect that in many intended target systems for CGD, such as mosquitoes, these conditions are typically exceeded. Estimates of single nucleotide mutation rates tend to be on the order of 10^−9^ – 10^−8^ in such species^18–20^ and a single mutation in the PAM motif suffices to create a resistance allele^10^. In addition, indels should occur within the 20 bp-long target sequence at rates comparable to the single nucleotide mutation rate^21^. We have shown that for values of *μ* in this range, resistance is already likely when *N_e_* = 10^6^, which is compatible with estimates from levels of neutral diversity in insect populations^22^. In many systems, the short-term values of *N_e_* relevant for a process as rapid as CGD may even be much larger than these values obtained from neutral diversity^17,23^, which will often be dominated by seasonal population crashes (e.g., caused by the winters in a temperate region) and historical bottlenecks^24,25^. Census population sizes in insects can easily reach billions and more at certain times of the year and resistance alleles will inevitably arise during these times when mutations are not limiting. If several generations occur between collapses, these alleles could reach sufficiently high frequencies prior to the next crash to persist^26^. Resistance probabilities may then actually depend on whether a driver is released in the spring or the fall. Note that our theoretical model can be easily extended to study such scenarios (SI section 3.4). Finally, even when assuming a comparatively small population, say *N_e_* = 10^5^, this still puts a strong bound on NHEJ rates (*δ* < *1/N_e_*) to prevent resistance from this mechanism. Experimental studies suggest that NHEJ rates are much higher even when steps have been taken to suppress it^5^. Of course, resistance from NHEJ may not be relevant in some scenarios^3^.

Our modeling framework allows us to evaluate and compare specific CGD strategies. One proposed strategy is to break an existing gene (e.g., a gene involved in insecticide resistance)^3^. In this case, NHEJ-repair could achieve the intended result — a broken copy of the gene. Since these alleles should carry similar costs as the driver, they are not likely to rise in frequency. However, resistance may still evolve if some mutations change the target sequence without actually breaking the gene, or if the ability to drive itself is already associated with some fitness costs, as will likely be the case if off-target effects of the driver have not been completely suppressed^27–29^.

Another proposed strategy is the insertion of a new gene (e.g., a gene that prevents mosquitoes from transmitting malaria^5^). If the target site is a non-functional region, NHEJ could then create cost-free resistance alleles, making resistance highly probable. To avoid this, Esvelt *et al.^3^* suggested targeting an essential gene for cleavage, where incorporation of the driver would rescue gene function, whereas NHEJ would likely knock out the gene. Note that in this case resistance may still evolve if NHEJ does not always completely knock out the gene. One strategy that could substantially reduce resistance potential would be engineering a driver with fitness costs that are completely dominant (SI section 3.2).

Though we have shown that resistance is almost a foregone conclusion in many standard scenarios of CGD, the lag time before resistance alleles actually become frequent may still allow for effective intervention strategies in the short term. Figure 4a shows that it takes over 100 generations until resistance alleles reach 50% frequency in our standard model, whereas the driver reaches >99% frequency in less than 30 generations. This lag time depends on the selective advantage of the resistance allele compared to the driver and can be yet much longer if driver costs are lower (Fig. 4b). For some applications, this may be good enough, as subsequent CGD constructs could be designed and released that specifically target resistance alleles, based on regular surveys of genetic variation in the target gene — a process that may be more effective than the use of multiple gRNAs from the outset^3^. Note that our theory also allows the assessment of strategies involving the purposeful release of resistance alleles for controlling a drive.

It is clear that there is a need for more detailed modeling of CGD on several fronts. For example, we assumed that resistance is a binary trait, yet resistance levels could depend on number, type, and location of mutations in the target site^10^. Furthermore, resistance alleles may be created if homology-directed repair inserts the driver construct, but introduces errors that prevent it from driving. We also limited our study to resistance at the target site, even though *trans* resistance might be common. CGD hosts may harbor natural variation for Cas9 expression levels or may produce peptides or RNA that silences the CRISPR machinery^30^. The possibility of all these other resistance mechanisms suggests that our estimates for the probability that resistance evolves are likely conservative. Our assumptions of a panmictic population of constant size could also be overly simplistic, especially since drivers might be specifically designed to reduce population sizes. In such cases, the lag time between the loss of the wildtype and the spread of resistance could be crucial. For example, a deleterious driver that quickly reaches 99% frequency in the population may be sufficient to cause extinction, even though resistance would ultimately evolve in a model with constant population size. In structured populations, CGD might even result in local extinction when a deleterious driver allele becomes fixed in a local subpopulation before it manages to spread into other parts of the population.

Throughout this analysis we remained purposefully agnostic to the potential benefits and risks of the release of this type of potent biological technology. We do, however, acknowledge the need for extensive discussion among scientists, policy makers, and the public before release takes place^3,31,32^. It is our hope that this work facilitates such discussion.

## Acknowledgments

We thank Anthony Long for suggesting the NHEJ resistance concern and Jackson Champer and members of the Messer and Clark labs for helpful discussions. This work was supported by National Institutes of Health (NIH) grants R01-GM064590 to A.G.C. and K99-GM114714 to R.L.U.

### Author Contributions

R.L.U., A.G.C. and P.W.M. conceived of the project, R.L.U. and P.W.M. carried out the analyses and wrote the manuscript.

### Competing Interests

The authors declare that they have no competing financial interests.

## Supplementary Information

### 1 Population genetic model of CGD

#### 1.1 Definition

Consider a locus with three types of alleles: wildtype (0), driver (*d*), and resistance (*r*). In our model, we define a resistance allele to be any allele that cannot drive and also cannot be converted into a functioning driver by the CGD machinery (*i.e.*, complete resistance). Let *x_0_*(*t*), *x_d_*(*t*), and *x_r_*(*t*) denote the frequencies of these alleles in generation *t*. We assume that the driver is initially introduced into the population in generation *t* = 0 at frequency *x_d_*(0). Let *c* specify the rate at which the driver successfully creates a double-strand break in a driver/wildtype heterozygote (the cleavage rate). Let *δ* specify the fraction of cases in which the repair of such breaks creates a resistance allele that can no longer be targeted by the gRNA – the typical outcome when repair occurs by NHEJ rather than HR. Let *μ* specify the rate at which resistance alleles arise by *de novo* mutation in wildtype alleles in the germline. This specifies an effective rate that includes all possible mutations that can create a resistance allele at the locus, such as single nucleotide mutations in the PAM motif or frameshifting indels within the target sequence. We normalize fitness such that wildtype homozygotes have a fitness *ω*_00_ = 1, whereas all other genotypes can carry arbitrary fitness cost, *ω_ij_* = 1 − *s_ij_* ≤ 1. We model a large, panmictic population of variance effective population size *N_e_* and census population size *N*.

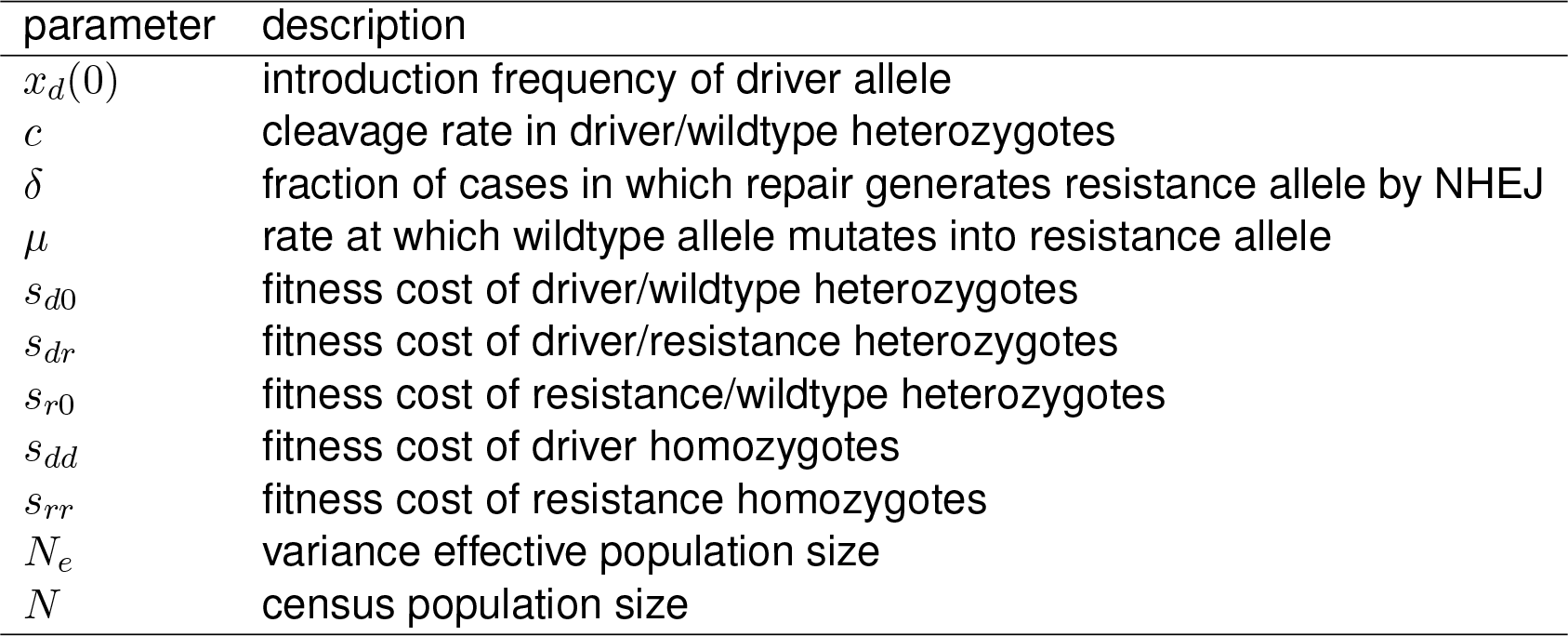

Note that generally there could also exist other alleles at the locus. For example, one may want to distinguish resistance alleles that arise by *de novo* mutation from those that arise by NHEJ, or resistance alleles that were already present as natural variation in the population. Depending on the specific drive scenario, these different resistance alleles could carry different costs. However, in our model we effectively combine all resistance alleles into a single class with the same fitness costs.

#### 1.2 Deterministic frequency dynamics

We initially assume that both *δ* and *μ* are small, such that the generation of new resistance alleles through *de novo* mutation or NHEJ does not noticeably affect wildtype and driver allele frequencies. Otherwise resistance will generally evolve quickly, as we will show below. Given allele frequencies *x_d_*, *x_r_*, and *x_0_* in generation *t*, the expected values of these frequencies in generation *t* + 1 are:

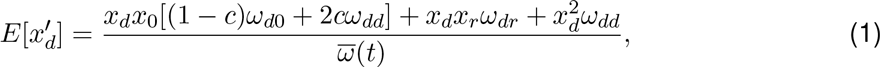

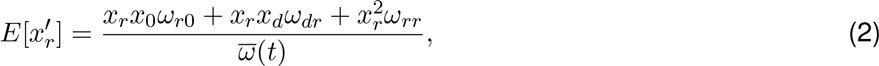

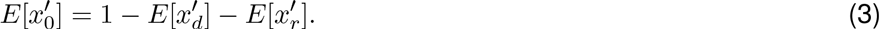

The contributions to the numerator in Equation (1) are as follows: The first term specifies the contributions of driver/wildtype heterozygotes that either successfully converted into driver homozygotes and thus now have fitness *ω_dd_*, which is expected to occur for a fraction *c*(1 − *δ*) ≈ *c* of driver/wildtype heterozygotes, or failed to convert and thus remain heterozygotes with fitness *ω_d0_*, which is expected to occur for a fraction 1 − *c*(1 − *δ*) ≈ 1 − *c* of driver/wildtype heterozygotes. The second and third terms specify the contributions from driver/resistance heterozygotes (fitness *ω_dr_*) and driver homozygotes (fitness *ω_dd_*), respectively. In Equation (2), the first term in the numerator specifies the contribution from resistance/wildtype heterozygotes, the second term specifies the contributions from resistance/driver heterozygotes, and the third term specifies the contribution from resistance homozygotes. Equation (3) directly follows from the fact that all three allele frequencies have to sum up to one.

The mean population fitness in generation *t* is given by:

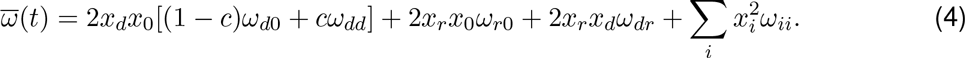

For any particular set of starting frequencies, these equations fully describe the population dynamics of the three alleles when assuming a deterministic scenario in which the alleles follow their expected allele frequencies exactly. This assumption will be appropriate when the alleles are segregating at high enough frequencies in a sufficiently large population, such that selective forces outweigh drift. However, we can also use these equations to specify expected values in Wright-Fisher-type simulations that explicitly incorporate random genetic drift.

#### 1.3 Driver dynamics in the absence of resistance

We and others have previously shown that these dynamics can already produce rich behavior in the absence of resistance (Unckless *etal*. 2015; Burt 2003). Possible outcomes can include the fixation of the driver, loss of the driver, and both stable or unstable equilibria, depending on the cleavage rate, *c*, the fitness costs of the driver allele, *s_dd_* and *s_d0_*, and the initial frequency *x_d_*(*t* = 0) at which the driver is introduced into the population. Briefly, in scenarios with high cleavage rate and small fitness costs, the driver will usually fix even when introduced at low frequency, whereas for low *c* and high fitness costs the driver may not be able to invade at all. Intermediate scenarios can give rise to equilibria that are usually unstable, but can be stable if *c* is small and fitness costs are low and recessive. For the remainder of this study, we will focus only on cases where, in the absence of resistance, the driver allele can go to fixation.

### 2 Resistance evolution

Resistance alleles can originate by three different mechanisms in our model: (i) they can already be segregating as standing genetic variation (SGV) in the population when the driver is introduced; (ii) they can arise by *de novo* mutation in wildtype genomes while the driver is spreading; and (iii) they can be created by the drive itself when cleavage repair via NHEJ results in mutated target sites that can no longer be recognized by the gRNA.

The overall likelihood that resistance evolves will depend on the supply of resistance alleles through the individual mechanisms, and the so-called establishment probability *π*(*t*) that a resistance allele successfully establishes in the population (i.e., is not lost by drift) when it is initially present in a single copy in generation *t*. Before we discuss the individual contributions of resistance from SGV, *de novo* mutation, and NHEJ, we will first show how establishment probabilities *π(t)* can be calculated in our model.

#### 2.1 Establishment probability of a single resistance allele

As long as resistance alleles are still rare, most of them should only be present in heterozygotes and we can therefore neglect resistance homozygotes. If a resistance allele is present at a very small frequency *x_r_*(*t*) in generation *t*, its expected frequency in the next generation will be:

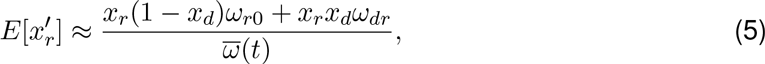

which can be rearranged into:

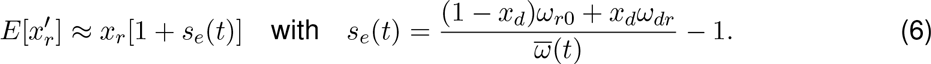

This ‘effective’ selection coefficient *s_e_*(*t*) specifies the expected change in frequency of the resistance allele in generation *t* under a model of exponential growth. Note that *s_e_*(*t*) is time-dependent, as it is a function of the driver frequency. If resistance alleles carry a cost themselves, then *s_e_*(*t*) could initially be negative and turn positive only after the driver has reached a certain frequency in the population.

To calculate *s_e_*(*t*) in a given generation *t*, we need to know the frequency *x_d_*(*t*) of the driver allele in that generation. Since we assume that resistance alleles are still at very low frequency, such that *x_d_*(*t*) ≈ 1 − *x_0_*(*t*), our dynamics from Equations (1)–(4) simplifies into:

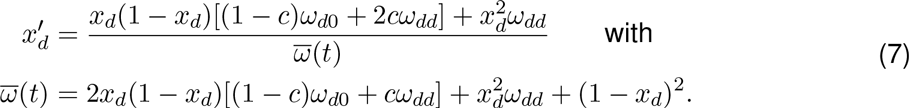

Assuming a deterministic model for the driver frequency, we can then calculate *x_d_(t)* recursively from its given starting frequency *x_d_*(*t* = 0).

Uecker and Hermisson (2011) recently derived the establishment probability of a new mutation for the general case that its selection-coefficient is time-dependent. Their theory can be directly adopted to our scenario if we assume that the new mutation is a resistance allele and its selection is given by *s_e_(t)*. According to Equation 16 in (Uecker and Hermisson 2011), the establishment probability of a resistance allele initially present in a single copy at time *t* is then given by:

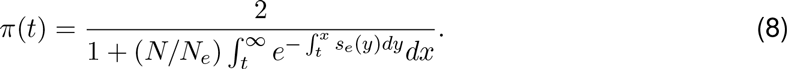

Note that the results from Uecker and Hermisson (2011) can also be used to model scenarios where census and effective population sizes are time-dependent (see section 3.4). Here we restrict ourselves to scenarios in which *N* and *N_e_* are constant.

In order to calculate the improper integral in the denominator of Equation (8) for *t* < *t*_fix_, we can split it into two components, prior to and after fixation:

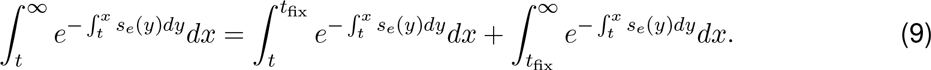

The integral in the exponent of the second summand still extends back to time *t*, but we can partition it as well into a component prior to *t*_fix_ and one afterwards, and then make use of the fact that *s_e_*(*t* ≥ *t*_fix_) = *s_e_*(*t*_fix_) will no longer depend on *t*. This yields:

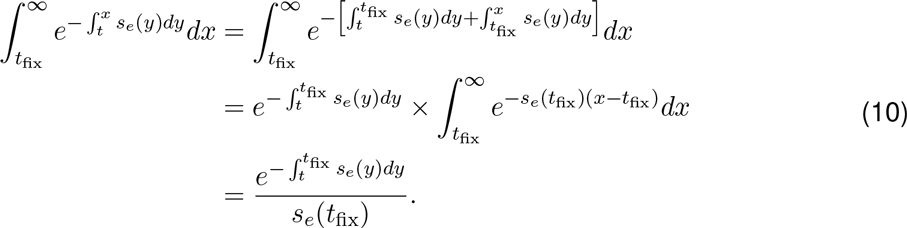

For *t* ≥ *t*_fix_, we simply obtain:

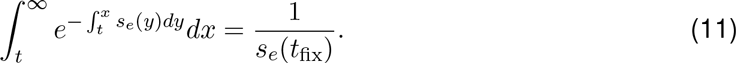

Both of these integrals diverge when *s_e_*(*t*_fix_) = 0, which could be the case if the fitness cost of the driver is completely dominant and resistance alleles would therefore not provide any fitness advantage in driver/resistance heterozygotes. We will discuss this particular scenario below in section 3.2. For now, we will assume that *s_e_*(*t*_fix_) > 0.

Importantly, all of the above integrals are defined over continuous time, whereas in our model generations are discrete and values for *s_e_*(*t*) are only defined for *t* ∈ ℕ_0_. Before we can estimate establishment probabilities in our discrete model, we first have to map these integrals onto sums over discrete generations. In the following, we will always use *i,t,t′* ∈ ℕ_0_ to denote discrete variables (generations), whereas *x,y* ∈ ℝ will denote continuous variables.

For mapping *s_e_*(*t*) onto continuous time, we will extend it to a piece-wise constant function with *s_e_*(*t* + *x*) = *s*(*t*) for 0 < *x* < 1. We can then associate the integrals inside the exponents in Equation (9), estimated between two discrete generations *t* < *t′*, with sums of the form:

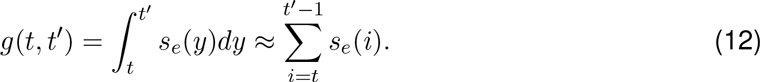

Discretization of the outer integral in Equation (9) can be achieved by partitioning it into the individual integrals between subsequent generations, yielding:

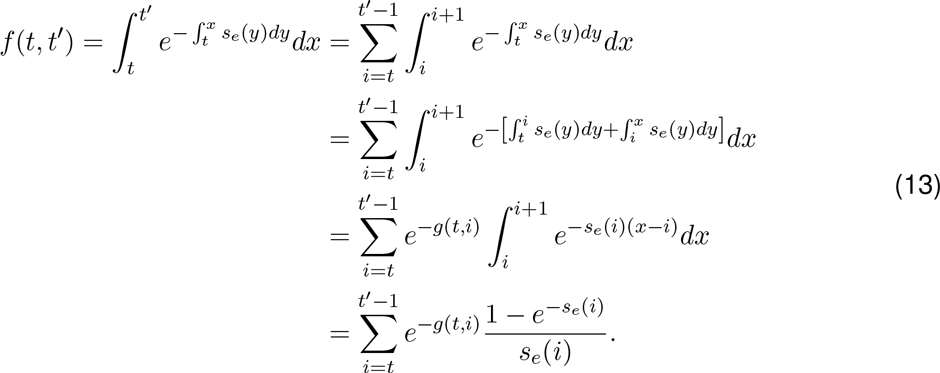

This way, both *g*(*t*, *t′*) and *f*(*t*, *t′*) are now successfully expressed in terms of only the values of *s_e_*(*i*) estimated in discrete generations *i* ∈ ℕ_0_. Combining all of the above results, we obtain for the establishment probability of a resistance allele arising in generation *t* in a single copy in our discrete model:

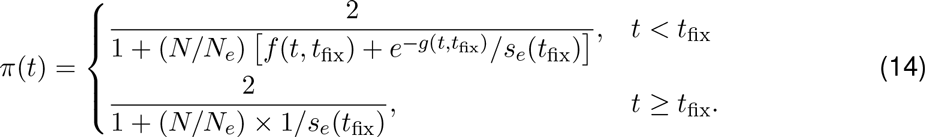

We can use this result to calculate the probability that a resistance allele will successfully establish in the population when it is initially present in *n_0_* copies at time *t*:

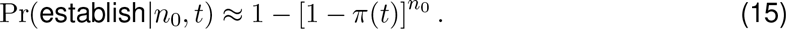

#### 2.2 Numerical analysis and standard model

Given that the number of parameters in our model is already quite large, we will typically perform our analyses using a ‘standard’ model, for which we then vary individual parameters (or pairs thereof) independently, while keeping the other parameters constant. This will allow us to dissect the particular role each individual parameter plays in the general behavior of the model, and the evolution of resistance specifically.

For this standard model, we assume a scenario in which cleavage is efficient, *c* = 0.9, as one would likely aim to design a driver this way. We further assume that resistance alleles carry no cost, *s_r0_* = *s_rr_* = 0, which we will vary later. Driver alleles, by contrast, carry a substantial cost that is codominant, *s_d0_* = 0.1 = *s_dd_*/2. In those analyses where both the driver and the resistance allele carry a cost, we assume that these costs are codominant to each other, *s_dr_* = *s_d0_* + *s_r0_*. We further set *δ* = 10^−6^ and *μ* = 10^−8^. Both are likely conservative choices with respect to the actual rate at which resistance alleles arise in many systems of interest, such as insects (see discussion in main manuscript). For simplicity, we assume *N* = *N_e_* = 10^6^. Driver alleles are introduced into the population at frequency *x_d_*(*t* = 0) = 10^−5^.

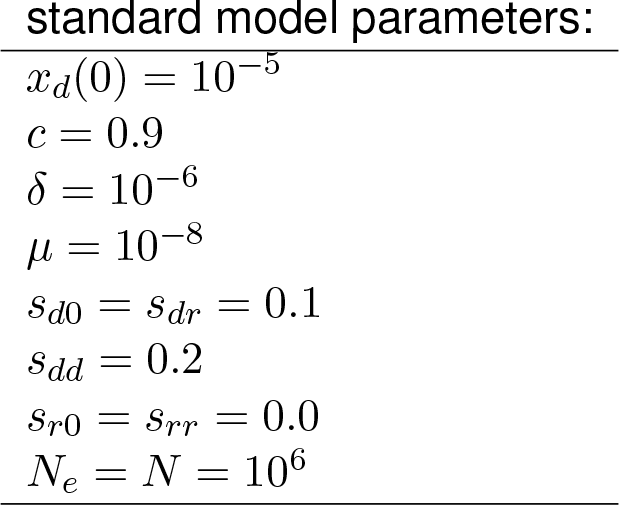

#### 2.3 Probability that resistance evolves from the SGV

Whether resistance against CGD can evolve from alleles that are already present in the SGV when the driver is introduced is a special case of the well-studied problem of evolutionary rescue from SGV (Orr and Unckless 2014). In particular, Hermisson and Pennings (2005) have previously developed a framework for calculating the probability that a population adapts to an environmental shift by utilizing alleles from the SGV that were previously neutral or deleterious, but became advantageous after the environmental shift. We can directly map this framework onto the evolution of resistance against a driver from the SGV. In this case, the environmental shift is the introduction of the driver, which can render a previously neutral or deleterious resistance allele beneficial as mean population fitness declines when the driver spreads in the population.

We assume that prior to introduction of the driver resistance alleles arise at rate *μ* per generation per haploid wildtype genome, and that they are evolving under mutation-selection-drift balance, specified by fitness costs *s_r0_* and *s_rr_* in heterozygotes and homozygotes, respectively. We define *θ* = 4*N_e_μ* to be twice the effective population-level mutation rate towards resistance alleles. After introduction of the driver, the fitness effects of resistance alleles are then given by *s_e_*(*t*) defined in Equation (6).

Let *P*_SGV_ denote the probability that resistance evolves from any allele present in the SGV at the time the driver is introduced, assuming that SGV is the only possible source of resistance alleles, which we can assure in our model by setting *μ* and *δ* to zero once the driver has been introduced. In this case, we obtain:

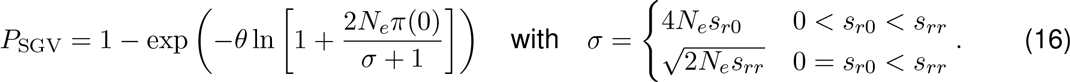

The codominant result (0 < *s_r0_* < *s_rr_*) follows directly from Hermisson and Pennings (2005). Specifically, *h′s_d_* in their Equation (8) specifies the heterozygous fitness cost of resistance alleles, which correspond to *s_r0_* in our model. The factor 2*hs_b_* in their equation is the establishment probability of a mutation present in a single copy in generation zero, which corresponds to *π*(0) in our model. The result for recessive resistance costs (0 = *s_r0_* < *s_rr_*) follows from the discussion provided after Equation (A11) in Hermisson and Pennings (2005), where they show that the factor 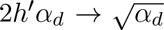 in their Equation (8) needs to be replaced by 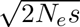 in the recessive case. The factor *s* in this expression specifies the homozygous fitness cost of the mutation, which corresponds to *s_rr_* in our model.

**Figure 1:**
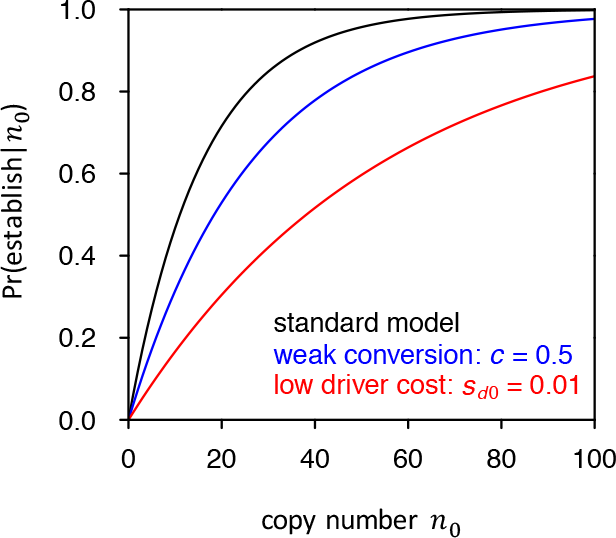
Probability that resistance establishes from the SGV in our standard model as a function of the initial number of copies (*n*_0_) in which the resistance allele is present when the driver is introduced. The black curve shows the prediction of Equation (15) for our standard model. The blue curve shows a scenario in which the driver cleaves at only 50% efficiency. The red curves shows a scenario where driver costs are ten times smaller than in our standard scenario.

Whether resistance evolves from SGV is primarily a function of the number of resistance alleles present when the driver is introduced (Figure 1). Lower cleavage rates slightly decrease establishment probabilities. This is because for a slower driver it will take longer until resistance alleles from the SGV experience their effective fitness advantage, increasing their chances of being lost to drift in the meantime. Similarly, lower driver costs will also decrease *P*_SGV_, as resistance alleles will have a smaller fitness advantage throughout.

Figure 2 shows *P*_SGV_ as a function of *θ* in our standard model, while simultaneously varying the cleavage rate (Figure 2a), the fitness cost of the driver allele (Figure 2b), or the fitness cost of resistance (Figure 2c,Figure 2d). Generally, as long as resistance alleles provide a net fitness advantage over the driver, evolution of resistance from the SGV is practically assured whenever *θ* ≥ 0.1, whereas it remains unlikely for *θ* ⪡ 0.1. This is consistent with the more general result from Hermisson and Pennings (2005), who showed that the probability of adaptation to an environmental shift using alleles from the SGV should depend only weakly on the selective advantage of these alleles in the new environment. Instead, it should be mostly determined by how many such alleles were present when the environment changed, which depends on *θ* and the selective disadvantage of these alleles prior to the change.

If resistance alleles carry a fitness cost themselves, this lowers the probability that resistances evolves from the SGV. However, this effect is much more pronounced when fitness costs are codominant than when they are recessive, consistent with the fact that even a deleterious allele can still reach a noticeable frequency in the SGV as long as its costs are only recessive. In our model, even for substantial recessive costs (*s_rr_* = 0.1), we still find *P*_S_gv to be practically indistinguishable from the cost-free scenario.

**Figure 2:**
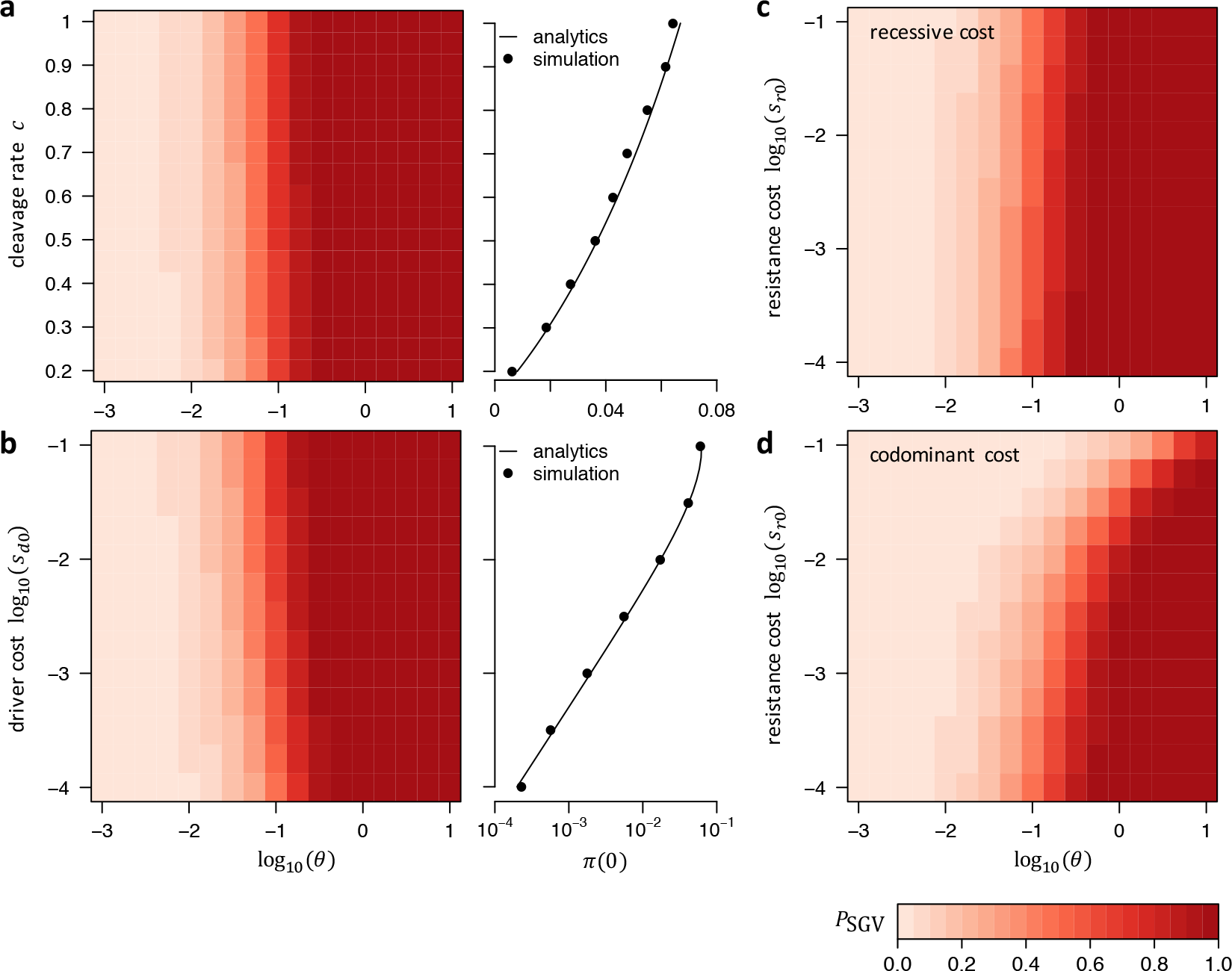
Probability that resistance evolves from SGV in our standard model. (**a**) *P*_SGV_ as a function of *θ* and cleavage rate. Higher *c* increases *P*_SGV_ by increasing *π*(0), because resistance alleles can experience their net fitness advantage faster when the driver spreads faster (panel to right). Dots show that our analytics agree well with numerical simulations under a Wright-Fisher model with conversion, selection, and drift. (**b**) *P*_SGV_ as a function of *θ* and driver cost. Higher driver costs also increases *P*_SGV_ by increasing *π*(0) (panel to right). Dots again show numerical simulations under a Wright-Fisher model. To vary *θ*, we always varied *N_e_* while keeping *μ* = 10^−8^ constant. Once *θ* becomes on the order of 0.1 or larger, establishment of resistance alleles from the SGV becomes generally likely. Higher cleavage rates, as well as higher fitness costs of the driver, both increase *P*_SGV_ by increasing the establishment probability *π*(0) of resistance alleles present at the time the driver is introduced (right panels). (**c**) *P*_SGV_ when resistance alleles carry a recessive fitness cost (*s_rr_* > *s_r0_* = 0). (**d**) Same as **c**, but assuming codominant fitness costs (*s_r0_* = *s_rr_*/2 > 0). Higher fitness cost of resistance alleles lead to lower *P*_SGV_ only if fitness costs are codominant, whereas recessive costs have almost no noticeable effect.

#### 2.4 Probability that resistance evolves from de novo mutation

In our model, resistance alleles can also be arise by *de novo* mutation after the driver has been introduced. We expect such alleles to be created in the population at rate:

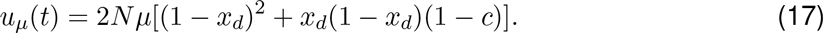

Any given resistance allele that arises as a single copy in generation *t* will successfully establish with probability *π(t)*, assuming that resistance does not arise from any other mechanism. Thus, the overall probability that at least one allele that arose prior to generation *t* establishes in the population is given by:

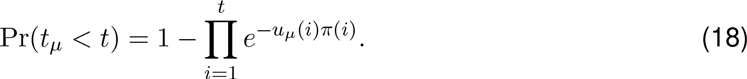

The factor e*^−u_μ_(i)π(i)^* here specifies the probability that no resistance allele from generation *i* successfully establishes in the population.

Note that *u_μ_(t)* is defined in terms of the census population size *N*, describing the actual number of individuals in the population. If the effective population size differs from the census size, this will not affect Equation (17), but will change the establishment probability of a new allele specified by Equation (8). However, this change is typically well-approximated by *π* ⟶ (*N_e_/N*)*π*, and *N* thus simply cancels out in the products *u_μ_(i)π(i)*. Note that in our numerical analyses we always set *N* = *N_e_* for simplicity.

Resistance alleles from de novo mutations can only arise as long as the driver has not yet fixed in the population. Afterwards, there will no longer be wildtype alleles present that could mutate into resistance alleles: therefore *u_μ_*(*t* ≥ *t*_fix_)= 0. The overall probability that resistance establishes from any *de novo* mutation arising during the drive is then:

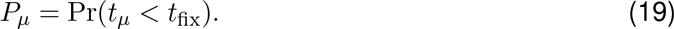

Figure 3 shows how *P_μ_*, depends on the parameters in our standard model. Again, we find that the key parameter determining the likelihood of resistance is *θ* = 4*N_e_μ*. However, in contrast to resistance from SGV, both the cleavage rate of the drive (*c*) and the cost of the driver allele (*s_d0_*) now noticeably affect *P_μ_*, as well. Specifically, lower cleavage rates increase *P_μ_* (the opposite effect they had on *P*_SGV_). This is because lower cleavage rates slow down the spread of the driver, providing more time for resistance alleles to emerge by *de novo* mutation before the driver becomes fixed. Lower fitness costs of the driver, on the other hand, decrease *P_μ_* by reducing the establishment probability of resistance alleles throughout the process, as resistance alleles will have a smaller net fitness advantage compared with the population mean.

If resistance alleles carry (codominant) fitness costs themselves, this has only a marginal effect on *P_μ_* as long as they still provide a net fitness advantage over the driver. Since we assume that resistance homozygotes are irrelevant for the establishment probability of a new resistance allele (see section 3.2 for a relaxation of this assumption), recessive costs have no noticeably affect on *P_μ_*, in our model.

When comparing *P*_SGV_ with *P_μ_* for the the same set of model parameters, we find that *P*_SGV_ is generally higher than *P_μ_*. Thus, resistance is more likely to evolve from resistance alleles already present when the driver is introduced, than those that arise from *de novo* mutation after its introduction. The only exception would be a scenario of a very inefficient drive (small *c*), in which case it would take a long time until the driver becomes prevalent in the population, reducing *π*(0) while at the same time providing more time for *de novo* alleles to emerge.

**Figure 3:**
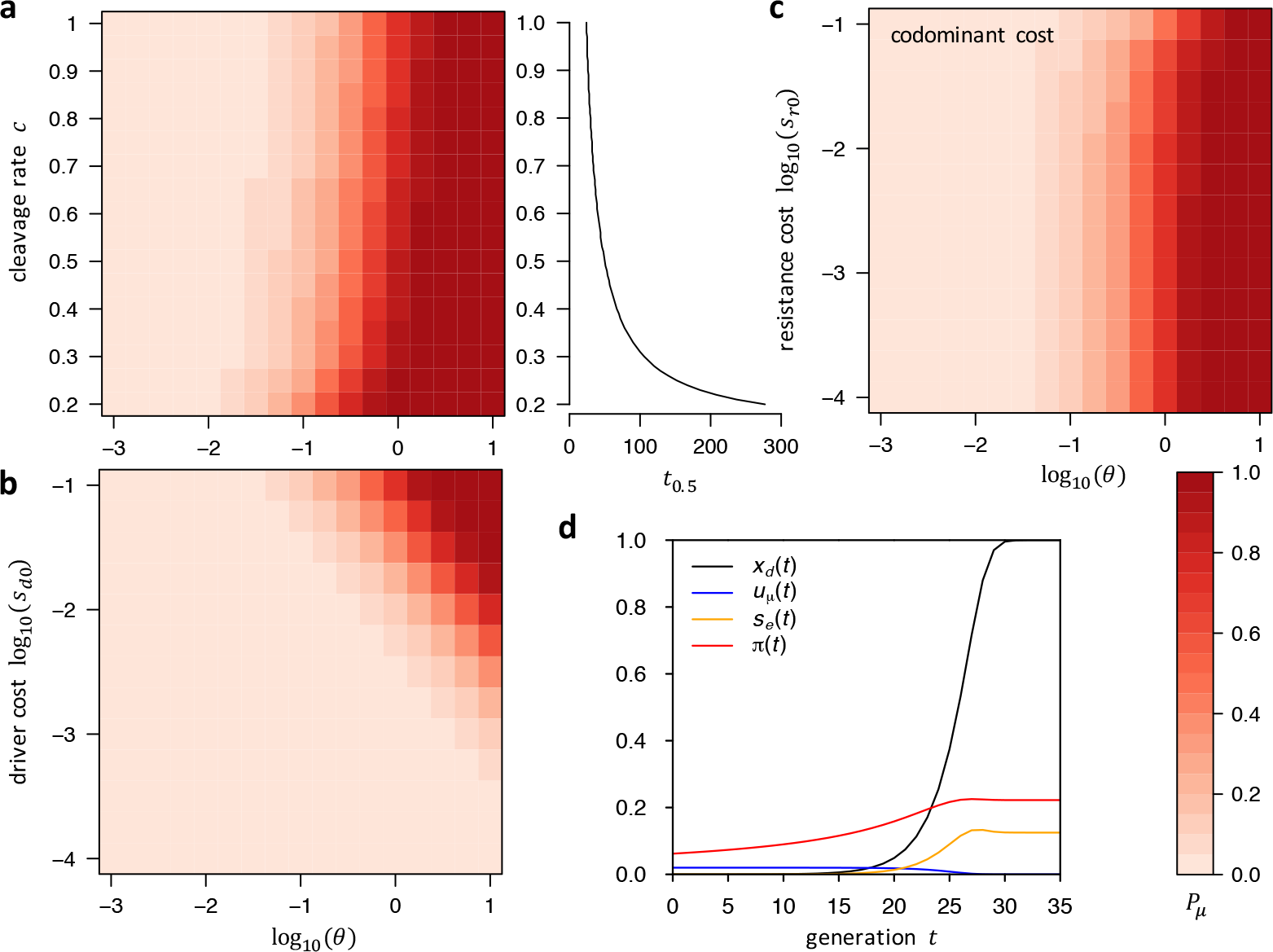
Probability that resistance evolves from *de novo* mutation in our standard model. (**a**) *P_μ_* as a function of *θ* and cleavage rate. The right panel shows how the waiting time until the driver reaches 50% frequency (*t*_0.5_) increases as cleavage rate decreases, which increases *P_μ_* by providing more time for resistance alleles to arise during the spread. (**b**) *P_μ_* as a function of *θ* and driver cost. Lower driver costs reduce *P_μ_* by lowering *π(t)* throughout the drive. (**c**) *P_μ_* as a function of *θ* and (codominant) resistance costs, which turn out to have only very little effect on *P_μ_*. To vary *θ* in **a**-**c**, we always varied *N_e_* while keeping *μ* = 10^−8^ constant. (**d**) Dependence between driver allele frequency *x_d_(t)*, overall rate *u_μ_*(*t*) at which resistance alleles arise by *de novo* mutation, their effective selective advantage *s_e_(t)* compared with the population mean, and the establishment probability *π(t)* of a resistance allele arising in a single copy in generation *t*. The driver allele fixes in approximately 35 generations in our standard model (unless resistance evolves). As the driver increases in frequency, so does *s_e_(t)*. The *de novo* mutation rate *u_μ_*(*t*) monotonically decreases as the driver displaces wildtype individuals from the population. The establishment probability *π(t)* also increases as *s_e_*(*t*) increases. Note, however, that *π*(*t* = 0) is already quite high in generation zero because even early resistance alleles have to survive drift for only a few generations before their relative selective advantage becomes noticeable in our model.

#### 2.5 Probability that resistance evolves from NHEJ

Lastly, even when both SGV and *de novo* fail at producing a resistance allele that successfully establishes in the population, resistance can still evolve from alleles created by the drive itself, when cleavage-repair by NHEJ results in a mutated target site. In our model, resistance alleles are created by this mechanism at rate:

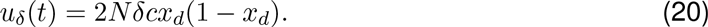

This rate is proportional to the number of driver/wildtype heterozygotes in the population and will thus be maximal when the driver is at intermediate frequency (Figure 4a). By comparison, the rate at which *de novo* resistance alleles arise is proportional to the overall number of wild-type alleles in the population, which is highest when the driver is introduced and decreases monotonically afterwards (Figure 3d).

As with *de novo* mutation, any given resistance allele that arises as a single copy in generation t by this mechanism will successfully establish with probability *π(t)*, assuming that resistance does not arise from any other mechanism. The overall probability that at least one allele that arose prior to generation *t* establishes in the population is given by:

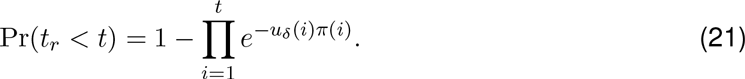

This equation is analogous to Equation (18) for the *de novo* mutation scenario, except for *u_μ_*(*t*) being exchanged by *uδ*(*t*). Given that *uδ*(*t*) will be zero once the the driver has fixed, we can again use this result to calculate the overall probability that at least one resistance allele from this mechanism establishes in the population:

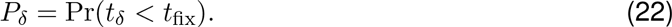

Figure 4 shows how *P_δ_* depends on the parameters in our standard model. To ensure that there is no resistance from the SGV or de novo mutation, we set *μ* = 0. We then varied *N_e_δ* by varying *N_e_* while keeping *δ* = 10^−6^ constant. As we discuss in the main text, this is likely to be a very low estimate for the rate at which cleavage repair generates resistance alleles through NHEJ in many systems, which may be rather difficult to achieve in practice. Higher rates will always increase *P_δ_*, making this estimate conservative with regard to the probability that resistance evolves by this mechanism.

Given the similarity between the NHEJ and de novo mutation scenarios, it is not too surprising that the key parameter determining the likelihood of resistance in the NHEJ scenario is the product *N_e_δ* similar to the parameter *θ* = 4*N_e_μ* in the de novo scenario (the factor 4 is not essential and primarily reflects a historical convention). Whenever *N_e_δ* becomes on the order of one or larger in our model, evolution of resistance from NHEJ becomes likely.

In contrast to the de novo mutation scenario, the cleavage rate has very little impact on *P_δ_*. This can be understood from the fact that the overall number of resistance alleles that arise by NHEJ is proportional to the overall number of cleavage events that occur throughout the spread of the driver, which does not depend strongly on *c* (as long as the spread of the driver is still dominated by conversion, rather than selection). For example, if drivers would not carry any fitness costs, fixation would, on average, require 2*N* conversion events throughout the process, regardless of whether the drive is fast or slow. Fitness costs of the driver, in contrast, do have a strong impact on *Pδ*, with higher fitness costs leading to lower *Pδ* by reducing *π(t)* throughout the process — the same effect they had in the *de novo* mutation scenario.

**Figure 4:**
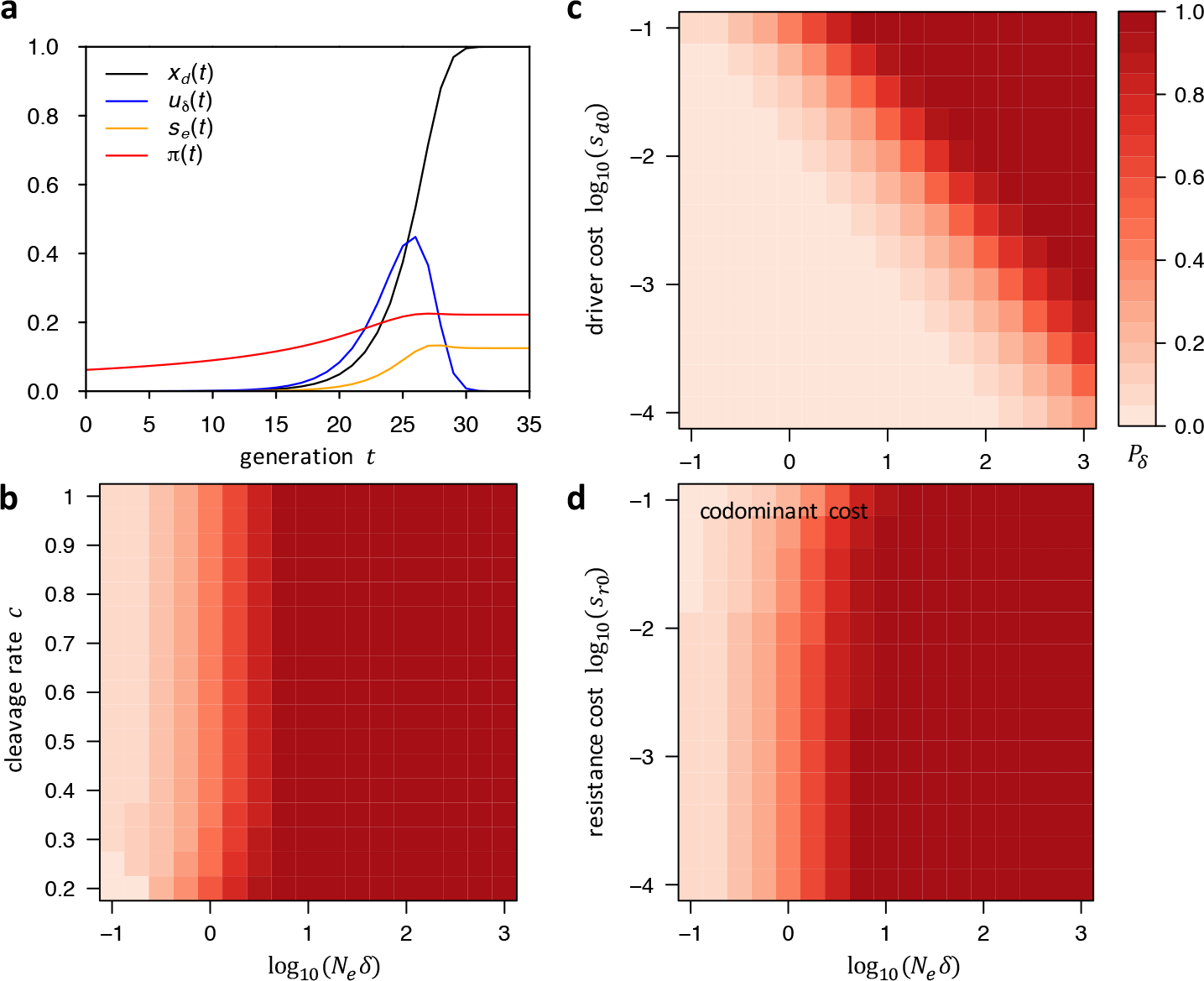
Probability that resistance evolves from NHEJ in our standard model. (**a**) Similar to Figure 3d but showing *uδ(t)* instead of *u_μ_(t)*. The overall rate at which resistance alleles are created by NHEJ is proportional to the overall number of driver/wildtype heterozygotes in the population, and thus maximal when the driver is at intermediate frequencies. (**b**) *P_δ_* as a function of *N_e_δ* and cleavage rate, which has very little effect on *P_δ_*. (**c**) *P_δ_* as a function of *N_e_δ* and driver cost, which lower *P_δ_* by reducing *π(t)*, similar to their impact on *P_μ_*. (**d**) *P_δ_* as a function of *N_e_δ* and (codominant) resistance costs, which have only very little effect on *P_δ_*. In **b**-**d** we varied *N_e_δ* by varying *N_e_* while keeping *δ* = 10^−6^ constant.

#### 2.6 Total probability of resistance

In the previous sections, we calculated the individual probabilities *P*_SGV_, *P_μ_* and *P_δ_*, that resistance evolves from each of the three mechanisms (SGV, *de novo* mutation, NHEJ, respectively), assuming that is has not evolved by any of the other mechanisms. We can directly combine these individual probabilities to calculate the total probability that resistance evolves by any of the three mechanisms:

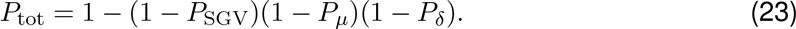

In our findings for the individual mechanisms, this total probability will depend primarily on three factors: *N_e_*, *μ*, and *δ*. Specifically, we have shown that whenever *θ* = 4*N_e_μ* becomes on the order of 0.1 or larger, resistance is likely to evolve from the SGV, as long as resistance alleles provide a net fitness advantage over the driver. This result was largely independent of the conversion rate of the driver, as well as the absolute fitness cost of the driver. Resistance alleles that arise from *de novo* mutation after introduction of the driver are generally less likely to contribute to resistance than alleles from the SGV. We further found that NHEJ is expected to produce resistance in the population whenever *N_e_δ* becomes on the order of 1 or larger, unless fitness costs of the driver are very small. Figure 5 summarizes these results by showing *P*_tot_ for a wide range of parameters and limiting cases in our standard model.

### 3 Special cases

#### 3.1 Frequent NHEJ regime

Our dynamical model for the allele frequencies described by Equations (1)–(4) assumed that the rates at which resistance alleles are created by mutation and NHEJ are small enough that we can neglect their actual contribution to changes in allele frequencies between generations. This may no longer be the case if NHEJ becomes more frequent, say *δ* = 0.01.

Extending our dynamical model such that we no longer neglect the contribution of NHEJ to changes in allele frequencies between generations is straightforward. In this case, the dynamics described by Equations (1)–(4) becomes:

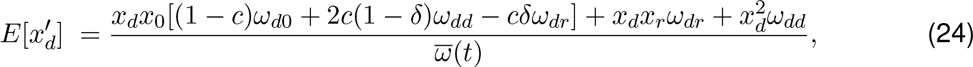

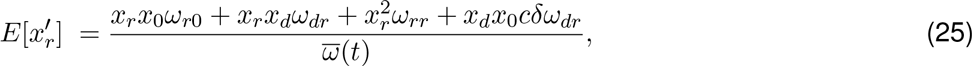

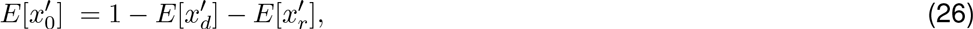

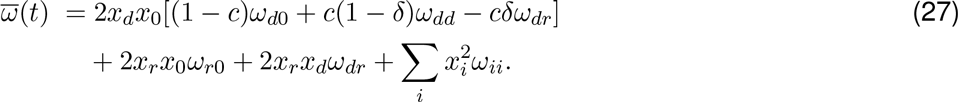

Evolution of resistance is practically inevitable in this regime as long as resistance alleles carry any fitness advantage relative to driver alleles, given that there will always be ample supply of resistance alleles being generated by NHEJ. In fact, if *δ* is sufficiently large, resistance alleles could become common even when driver alleles carry no cost at all. In this case, if the driver is initially introduced at very low frequency, the ratio of resistance allele frequency over resistance allele frequency should simply be:

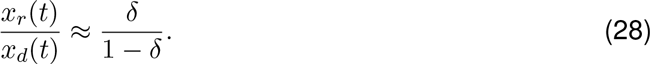

**Figure 5:**
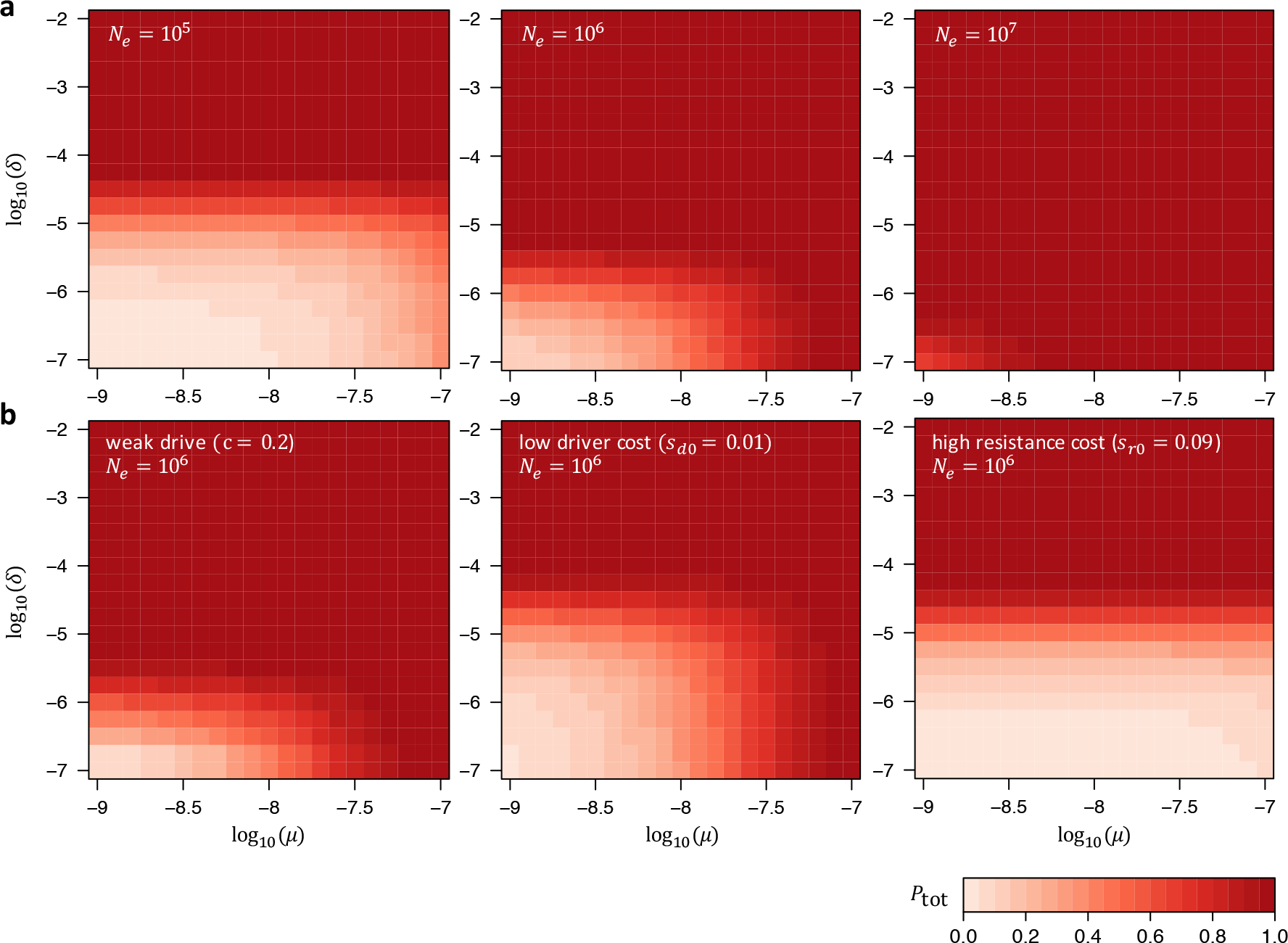
Total probability that resistance evolves in our standard model. (a) *P*_tot_ as a function of *μ* and *δ* for three different effective population sizes *N_e_* = 10^5^,10^6^,10^7^. Larger values of *N_e_* increase the probability of resistance, yet even in the scenario with *N_e_* = 10^5^ low values of *δ* < 10^−5^ are still required if resistance from NHEJ is to be prevented. (b) *P*_tot_ as a function of *μ* and *δ* for fixed *N_e_* = 10^6^ in three limiting cases. Left: scenario of a very weak drive with cleavage rate: *c* = 0.2. Comparison with the center panel in a shows that this has almost no effect on *P*_tot_. Center: scenario with low driver costs, *s_d0_* = *s_dd_*/2 = 0.01. While this allows for larger values of *δ*, the dependence on *μ* does not change much, since *P*_S_gv remains largely unaffected by the driver costs (compare with Figure 2b). Right: scenario in which resistance alleles carry (codominant) fitness costs, *s_r0_* = 0.9*s_d0_* = *s_rr_*/2 = 0.09, that are almost as large as the costs of the driver. In this case, resistance is unlikely to evolve from SGV or *de novo* mutation. However, it is still likely to evolve by NHEJ unless *δ* < 10^−5^.

This is because every time CGD-induced cleavage in a driver/wildtype heterozygote is repaired, in a fraction *δ* of cases a resistance allele will be added to the population, whereas in a faction 1 − *δ* of cases a diver allele will be added.

Fitness differences between driver and resistance alleles will change their relative frequencies over time, but this can take quite some time unless these differences are very large. For a fast drive in the large *δ* regime, we would expect that at the time the wildtype allele is lost from the population, the relative frequency of resistance alleles should still be close to the ratio *δ*/(1 − *δ*) at which these alleles were originally created in the population.

#### 3.2 Recessive and dominant driver costs

So far we have assumed that fitness costs of driver alleles are codominant. If those fitness costs were recessive, the driver would be able to initially rise in frequency faster, as driver/wildtype heterozyogtes would carry no cost. Conversely, if fitness costs were dominant, this would slow down the initial rise of the driver.

However, in our standard model the expected differences in driver frequency trajectories between scenarios with recessive, codominant, and dominant fitness costs are only marginal (main manuscript Figure 2a). In our standard model, frequency changes of the driver allele between generations are still dominated by conversion of heterozygotes into homozygotes, whereas selection would only become important once driver fitness costs would actually be of the same order as the cleavage rate, *s_d0_* ≈ *c*.

Dominance of the driver can nevertheless have strong impact on the probability that resistance arises, due to its effects on the fitness of driver/resistance heterozygotes. Once a driver is frequent, while resistance alleles are still rare, these alleles will predominantly be present in such driver/resistance heterozygotes. If driver costs are recessive, *s_e_*(*t*) will then be larger than in the codominant case, increasing *π*(*t*) and thus *P_μ_* and *P_δ_* (main manuscript Figure 2a).

By contrast, if driver costs are completely dominant, driver/resistance heteroygotes will have no fitness advantage over driver homozygotes. Once wildtype alleles have been completely displaced by the driver, only resistance homozygotes would then have a fitness advantage. In this case, our approach for calculating establishment probabilities based on Equation (14) breaks down, because it does not take resistance homozygotes into account.

However, resistance could still evolve in this scenario if enough resistance homozygotes are present at some time in the population, as they still have a fitness advantage. The question of how likely resistance is to evolve in this case is analogous to problem of whether a beneficial mutation can establish in a population when its fitness effects are completely recessive. Such a recessive mutation with fitness 1 + *s′* in homozygotes and fitness 1 in heterozygotes has an establishment probability of approximately 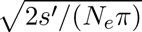 when initially present in a single copy (Kimura 1962), which we can extend to the probability that the mutation will successfully establish when it is initially present at any given frequency *x*_0_:

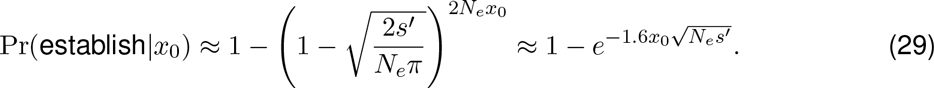

In our standard scenario with cost-free resistance, the net fitness advantage of resistance homozygotes in this case will be *s′* ≈ *s_dd_*/(1 − *s_dd_*), assuming that the driver is completely dominant and close to fixation. Equation (29) then allows us to calculate the establishment probability of a resistance allele present at frequency *x*_0_ when most individuals in the population already carry a driver allele. In order for resistance evolution to become likely in this scenario, the resistance allele has to be present at substantial initial frequency in the population. For example, when the driver has a dominant fitness cost of *s_dd_* = 0.1, resistance alleles would have to be present at ≈ 0.1% frequency in order to have a 50% chance of successfully establishing (main manuscript Figure 2b). Note that according to Equation (28) this is what we would in fact expect in scenarios with *δ* > 10^−3^.

#### 3.3 High introduction frequency

In our standard model, the driver is introduced at frequency *x_d_*(0) = 10^−5^ in generation zero (corresponding to 20 copies in a diploid population of census size *N* = 10^6^). We have not varied this introduction frequency in our previous analyses, so one might wonder whether higher or lower frequencies can affect the probability that resistance evolves.

Figure 6 shows that changing *x_d_*(0) has almost no noticeable effect on the total probability that resistance evolves. This is the consequence of two competing effects: A higher introduction frequency will increase *π*(0) and thus *P*_SGV_, compared with a lower introduction frequency, as resistance alleles present in the SGV will experience their fitness advantage faster. However, a higher introduction frequency will also lead to faster fixation of the driver, thus leaving less time for *de novo* resistance mutations to occur, which will decrease *P*_SGV_ The two effects approximately cancel out. Furthermore, *P_δ_* will generally not depend much on the introduction frequency (at least as long as *x_d_*(0) is still much smaller than one), because almost all of the resistance alleles produced by NHEJ arise when the driver is at intermediate frequency.

Note that changing *x_d_*(0) will nevertheless have a noticeable impact on the time it takes for the driver to become frequent in the population. For example, while it takes the driver *t*_0.5_ ≈ 26 generations to reach 50% frequency in our standard model with *x_d_*(0) = 10^−5^, it will only take *t_0.5_* ≈ 10 generations when the driver is introduced at frequency *x*(0) = 0.01.

#### 3.4 Varying population size

We have so far assumed that population size remains constant over time. This assumption is likely violated in many systems. Insect populations, for example, can often show dramatic fluctuations in population size over time scales of just a few generations (Berryman 2002). In temperate areas, winters can result in population crashes, followed by rapid increase during the growing season. Management practices for pests, such as pesticide application, can also influence population dynamics substantially. Finally, CGD itself could affect population size if the goal is to spread a harmful allele in the population.

Relaxing the assumption of constant population size can impact three aspects of our analytical framework for calculating resistance probabilities: (i) the supply of resistance alleles in the SGV, (ii) the rates *u_μ_(t)* and *u_δ_(t)* at which resistance alleles arise by *de novo* mutation and NHEJ, and (iii) the establishment probability *π(t)* of a resistance allele initially present in generation *t* in a single copy.

Fortunately, the theoretical result by Uecker and Hermisson (2011) we adopted for calculating *π(t)* in our CGD model does allow for arbitrary changes in population size over time. We just used a simplified version of the result for the special case of constant population size. In the general case, given functions *N*(*t*) and *N_e_*(*t*) for census and variance effective population sizes, Equation (8) becomes:

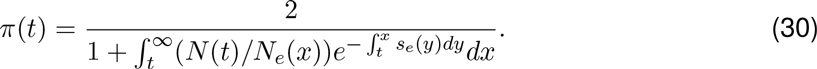

This equation can then be solved numerically for the given demography in the same way we did for the constant population size scenario.

**Figure 6:**
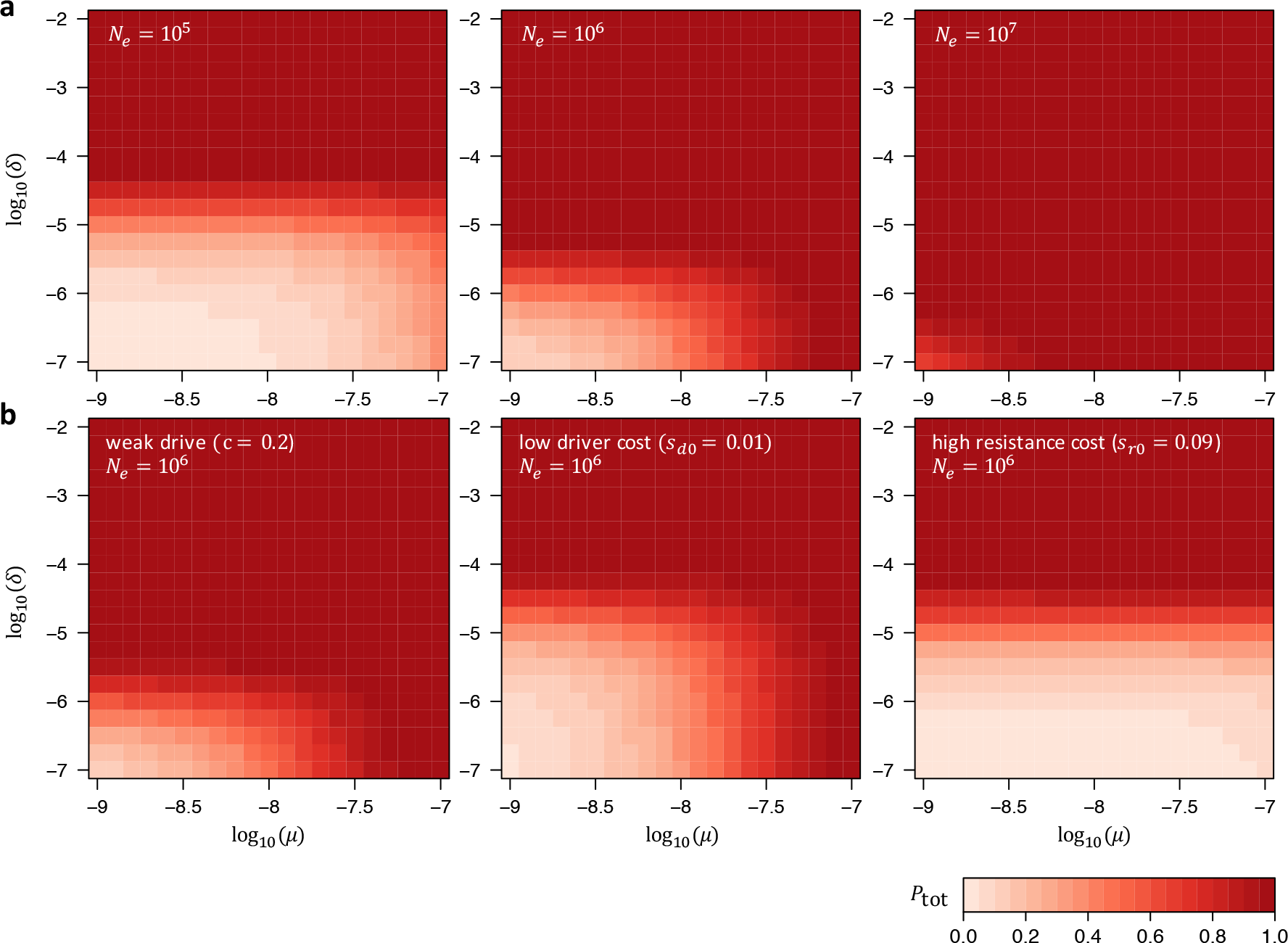
Total probability that resistance evolves in our standard model when the driver is introduced at frequency *x_d_*(0) = 0.01, instead of *x_d_*(0) = 10^−5^. Otherwise the plot is analogous to Figure 5. There are almost no noticeable differences between the two introduction frequencies.

Both *u_μ_*(*t*) and *uδ*(*t*) are already functions of *t* in our model. Under varying population size, we simply have to express these rates in terms of the census population size *N(t)*:

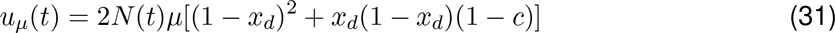

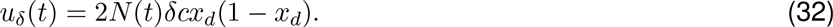

Given *π(t)* and rates *u_μ_*(*t*) and *uδ*(*t*), the resistance probabilities *P_μ_* and *P_δ_* can then be calculated according to Equations (19) and (22) the same way we did for the constant population size scenario.

To our knowledge, there are no general analytical solutions for mutation-selection-drift balance that allow for calculation of allele frequency distributions under arbitrary demography scenarios. Calculation of *P*_SGV_ will therefore typically rely on numerical simulations to infer the frequency of resistance alleles in the SGV. However, such distributions could be easily obtained from Wright-Fisher simulations under any given demographic model. In combination with *P*_SGV_, this will then allow numerical estimation of *P*_SGV_.

### 4 Software

Our numerical analyses are implemented in a C++ command line program. This program takes as input all parameters of the given model: *x_d_*(*t* = 0), *c*, *μ*, *δ*, *s_d0_*, *s_dr_*, *s_r0_*, *s_rr_*, *N_e_*, *N*. For these parameters, it then calculates, the driver allele frequency trajectory *x_d_*(*t*) under a deterministic model, specified by Equations (1)–(4) in the absence of resistance, the rate *u_μ_(t)* at which resistance alleles are expected to arise by *de novo* mutation, the rate *uδ*(*t*) at which resistance alleles are expected to arise by NHEJ, the effective selective advantage *s_e_*(*t*) of a resistance allele compared with the population mean, and the establishment probability *π(t)* of a resistance allele arising in a single copy in generation *t*. Results are provided for each generation 0 ≤ *t* ≤ *t*_fix_. The program also calculates the individual resistance probabilities *P*_tot_, *P*_SGV_, *P_μ_*, and *P_δ_*. Executables, source-code, and documentation for this program are available from the corresponding author upon request and will be made publicly available after acceptance of the manuscript.

